# Seasonal Dynamics in the Number and Composition of Coliform Bacteria in Drinking Water Reservoirs

**DOI:** 10.1101/2021.02.16.428560

**Authors:** Carolin Reitter, Heike Petzoldt, Andreas Korth, Felix Schwab, Claudia Stange, Beate Hambsch, Andreas Tiehm, Ilias Lagkouvardos, Johannes Gescher, Michael Hügler

**Affiliations:** TZW: DVGW-Technologiezentrum Wasser (German Water Center), Karlsruher Str. 84, 76139, Karlsruhe, Germany; TZW: DVGW-Technologiezentrum Wasser (German Water Center), Wasserwerkstr. 2. 01326, Dresden, Germany; ZIEL - Institute for Food & Health, Technical University of Munich, Freising, Germany.; Department of Applied Biology, Institute for Applied Biosciences, Karlsruhe Institute of Technology (KIT), Fritz-Haber-Weg 2, 76131, Karlsruhe, Germany.; Institute for Biological Interfaces, Biomolecular Micro- and Nanostructures, Karlsruhe Institute of Technology, Eggenstein-Leopoldshafen, Germany

**Keywords:** coliform bacteria, drinking water reservoir, mass proliferation, climate change, *Enterobacter*, *Lelliottia*

## Abstract

Worldwide, surface waters like lakes and reservoirs are one of the major sources for drinking water production, especially in regions with water scarcity. In the last decades, they have undergone significant changes due to climate change. This includes not only an increase of the water temperature but also microbiological changes. In recent years, increased numbers of coliform bacteria have been observed in these surface waters. In our monitoring study we analyzed two drinking water reservoirs (Klingenberg and Kleine Kinzig Reservoir) over a two-year period in 2018 and 2019. We detected high numbers of coliform bacteria up to 2.4 x 10^4^ bacteria per 100 ml during summer months, representing an increase of four orders of magnitude compared to winter. Diversity decreased to one or two species that dominated the entire water body, namely *Enterobacter asburiae* and *Lelliottia* spp., depending on the reservoir. Interestingly, the same, very closely related strains have been found in several reservoirs from different regions. Fecal indicator bacteria *Escherichia coli* and enterococci could only be detected in low concentrations. Furthermore, fecal marker genes were not detected in the reservoir, indicating that high concentrations of coliform bacteria were not due to fecal contamination. Microbial community revealed *Frankiales* and *Burkholderiales* as dominant orders. *Enterobacterales,* however, only had a frequency of 0.04% within the microbial community, which is not significantly affected by the extreme change in coliform bacteria number. Redundancy analysis revealed water temperature, oxygen as well as nutrients and metals (phosphate, manganese) as factors affecting the dominant species. We conclude that this sudden increase of coliform bacteria is an autochthonic process that can be considered as a mass proliferation or “coliform bloom” within the reservoir. It is correlated to higher water temperatures in summer and is therefore expected to occur more frequently in the near future, challenging drinking water production.

**Highlights:**

- Coliform bacteria proliferate in drinking water reservoirs *to values above* 10^4^ per 100 ml
- The genera *Lelliottia* and *Enterobacter can form these “coliform blooms”*
- Mass proliferation is an autochthonic process, not related to fecal contaminations
- It is related to water temperature and appears mainly in summer
- It is expected to occur more often in future due to climate change

**Graphical abstract:** 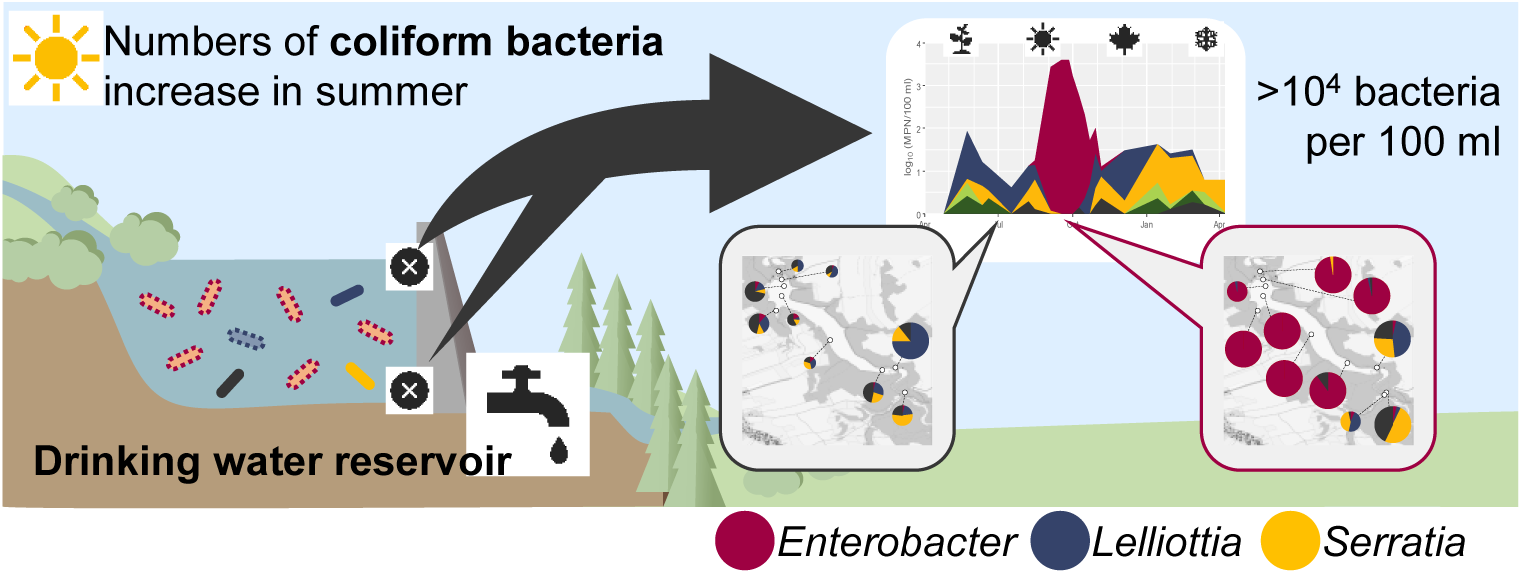

## 1. Introduction

Worldwide, surface water is a major source for drinking water production, estimated to cover approx. 50% of the worldwide need (WHO, 2016). Reservoirs or dams are built to store surface water to guarantee the availability of water for drinking water production, especially in regions with water scarcity. In the following, we refer to such dams and reservoirs used for the production of drinking water as “drinking water reservoirs”. In Germany about 12% of drinking water supply is covered by the use of surface water from lakes and reservoir waterworks (Statistisches Bundesamt, 2018). Regionally, this proportion is much higher, like e.g. in Saxony and Thuringia with about 50%.

Reservoirs are created by damming a river in a valley, usually with a pre-reservoir to withhold the coarsest dirt. From a limnologic point of view, reservoirs can be regarded as a mixture of lake and river. The closer to the dam, the more lacustrine the reservoir gets. In comparison to many lakes, the water retention time is considerably shorter. Generally, stratification is observed in summer. Temperature uniformity on the one hand and wind on the other hand ensure a mixing in spring and autumn. In winter, inverse stratification can occur. Stratification influences the distribution of substances in the reservoir, as mixing of water bodies no longer occurs. As a consequence oxygen consumption in the hypolimnion can lead to the solution of metals (e.g. manganese) and nutrients from the sediment.

Within the last decades, reservoirs have undergone significant changes due to climate change. One of the main factors is the temperature. In lakes, water temperature has increased around 0.1 to 1.5 °C in the last 40 years (Bates et al., 2008). Even the cold hypolimnetic water shows an increase in temperature of about 0.1 to 0.2 °C per decade (Dokulil et al., 2006). Temperature influences many physicochemical parameters and globally leads to an increase in pH and a decrease of dissolved gases like oxygen in deeper layers (Bates et al., 2008; Delpla et al., 2009). Furthermore, due to global warming, the stratification period in lakes in the Northern Hemisphere has lengthened by 2 to 3 weeks and thermal stability has increased (Bates et al., 2008) due to an earlier warming-up of the water and a decreased ice coverage in winter (Magnuson et al., 2000).

Unlike groundwater, that is protected by overlying soils, surface waters are vulnerable to various contaminations and often show microbial fecal contamination that impacts drinking water production or recreational activities. Rivers and streams are widely used as receiving waters for wastewater treatment plant effluents, and rivers, as well as lakes and reservoirs, are influenced by diffuse pollution like e.g. agricultural runoffs (Kirschner et al., 2017; Kistemann et al., 2002). Fecal pollution poses potential health risks to humans, therefore microbiological monitoring is essential in assessing the water quality of reservoirs used for drinking water production (Kistemann et al., 2002; Paruch et al., 2019).

The hygienic-microbiological water quality of drinking water, raw water for drinking water production and recreational water is examined using bacterial indicators such as coliform bacteria, *Escherichia coli* (*E. coli*) and enterococci (European Community, 1998). Especially the detection of the latter two indicates a possible fecal contamination. This is a serious health issue as it indicates the potential presence of pathogens. For coliform bacteria, this concept has been relativized as this heterogeneous group of *Enterobacteriacaea* also occurs in the environment, like e.g. water, plants and soil, thus their hygienic relevance is still under debate (Ashbolt et al., 2001; Leclerc et al., 2001; Octavia and Lan, 2014; Stevens et al., 2003; WHO, 2017).

In order to be able to operate reservoir water treatment in a natural, stable and economical way, it is necessary to provide raw water, which means the water used for drinking water production, with the lowest possible chemical and microbiological contamination. For this purpose, appropriate protection and management concepts for the catchment areas and raw water resources are in place. Despite the implementation of the proven management concepts, changes in the raw water quality occur, which pose considerable challenges for the treatment process.

In recent years, high densities of coliform bacteria have repeatedly been observed in drinking water reservoirs and lakes (Davis et al., 2005; Exner et al., 2005; Freier et al., 2005; Packroff and Clasen, 2005). The observed high concentrations of coliform bacteria (occasionally above 10^4^ per 100 ml) led to uncertainty for water suppliers and represented a challenge for drinking water treatment. Water works treating reservoir water often rely on classical treatment technologies like flocculation and filtration and final disinfection with chlorine or chlorine dioxide. According to the German recommendations, treated water should be free from indicator bacteria prior to final disinfection. With very high concentrations of coliform bacteria in the raw water, these requirements cannot always be fulfilled.

High numbers of coliform bacteria occur mainly in summer, which means at a time when high water temperatures prevail. As possible factors, besides water temperature, dissolved organic carbon (DOC) and oxygen content have been discussed (Exner et al., 2005; Freier et al., 2005; Kämpfer et al., 2008; Packroff and Clasen, 2005). In order to clarify the question of the cause of high concentrations of coliform bacteria, it is important to understand the ecology in a reservoir and the changes they have undergone in recent years due to climate change.

Therefore, the aim of our study was to examine the phenomenon of the “coliform blooms” in drinking water reservoirs in Germany. For this reason, microbial indicator bacteria like coliform bacteria, *E. coli* and enterococci that occur in drinking water reservoirs were quantified and afterwards identified using Matrix-assisted laser desorption ionization-time of flight mass spectrometry (MALDI-TOF MS) and Multilocus sequence analysis (MLSA). Especially *E. coli* and enterococci as fecal indicator bacteria (FIB) indicate a potential fecal contamination, which may involve a risk to public health. Microbial source tracking (MST) methods were applied in order to identify the origin of fecal contaminations. In addition, total cell counts (TCC) and the microbiome were analyzed, to view the results in context with the whole microbial community.

## 2. Material and methods

### 2.1. Study side and sampling

Sampling was carried out between April 2018 and December 2019 in two drinking water reservoirs in Germany (Klingenberg Reservoir in Saxony and Kleine Kinzig Reservoir in Baden-Württemberg). The main characteristics of the reservoirs are summarized in Table 1.

**Table 1:** General description of the Klingenberg Reservoir and Kleine Kinzig Reservoir analyzed during this study.

|  | Klingenberg Reservoir (R1) | Kleine Kinzig Reservoir (R2) |
| --- | --- | --- |
| State | Saxony | Baden-Württemberg |
| Location | Eastern Ore Mountains<br>(Erzgebirge) | Black Forest |
| River basin | Elbe | Rhine |
| Altitude | 390 m ASL | 605 m ASL |
| Ground level | 359.7 m ASL | 544 m ASL |
| Withdrawal points | 360.2 - 385 m ASL | 544 - 599.3 m ASL |
| Raw water | 360.2 m ASL | 564 - 564 m ASL |
| Surface water | 385 m ASL | 589.3 - 599.3 m ASL |
| Depth (Average) | 15 m | 22 m |
| Depth (Maximum) | 33 m | 65 m |
| Water surface | 116 ha | 56 ha |
| Capacity | 16 Mio m<sup>3</sup> | 13 Mio m<sup>3</sup> |
| Raw water delivery | 1000 l/s | 200 l/s |
| Retention time | 120 d | 190 d |
| Catchment area | 90 km<sup>2</sup> (forest and agriculture) | 18 km<sup>2</sup> (mainly forest) |
| Ecology | mesotrophic | oligotrophic |
| Geology | gneiss | fretted sandstone |

For the monitoring study, the surface water and raw water of the reservoirs was sampled on a two to four weeks basis. For Kleine Kinzig Reservoir sampling was carried out at a tower located 151 m away from the dam wall in the lake. This tower consist of seven sampling points for the measurement of the depth profile between ground level (544 m above sea level (ASL)) and the surface 55 m higher (599.3 m ASL). The withdrawal height of the raw water varies according to the seasons and physical conditions. During measuring period the withdrawal height was between 20 and 30 m above ground. Only in January 2019 the withdrawal height was higher (45.3 m above ground) due to high turbidity contents in the deeper water. Surface water was sampled in the highest possible withdrawal floor, which differed due to the water level between 45.3 m to 55.3 m above ground.

In Klingenberg Reservoir sampling spots are directly at the dam wall and consist of six sampling steps from nearly ground level (360.2 m ASL) up to the surface with 25.3 m (385 m ASL). The extraction height of raw water was 50 cm above ground throughout the entire series of measurements. Surface water was extracted at the highest withdrawal point 25.3 m above ground. For Klingenberg Reservoir it was also possible to monitor the main inflow (Wilde Weißeritz). With these water samples microbiological analysis and identification of bacteria were conducted as described in the next section.

In order to answer the question for parameters that influence the variability of coliform bacteria within the reservoir, a sampling campaign was carried out at Klingenberg Reservoir in June and September 2018, during which surface water of different points in the reservoir, the depth profile and all inflows were sampled as described below. In addition, 2 L water samples were collected from selected sites to investigate fecal markers as well as the microbial community. For the Kleine Kinzig Reservoir, water samples for fecal markers and microbial community were taken during sampling campaigns in July and December 2019 from the depth profile. These samples were pretreated and DNA was extracted as described below.

High concentrations of coliform bacteria have been observed in many drinking water reservoirs and lakes in Germany. Therefore, in addition to the two reservoirs intensively sampled in this study, coliform bacteria were also quantified and identified from other reservoirs and lakes when high densities occurred. In these cases, the responsible water suppliers provided either water samples or isolates. Those samples were treated as described later and isolates were identified using MALDI-TOF MS and MLSA (see below). The following surface waters were examined: Lake Constance, Rappbode Reservoir, Breitenbach Reservoir and Söse Reservoir.

Meteorological data (air temperature, precipitation, wind speed) were measured during the complete period of monitoring on a daily basis. Furthermore, physical and chemical parameters were documented according to the German regulations for water, sewage, and sludge analysis (Wasserchemische Gesellschaft and Deutsches Institut für Normung e.V., 2020). For measurement of on-site parameters (oxygen, temperature, conductivity) multiparameter probes were used (SEBA Hydrometrie GmbH & Co. KG, Kaufbeuren, Germany, at Kleine Kinzig Reservoir and Sea&Sun Technology GmbH, Trappenkamp, Germany, at Klingenberg Reservoir). For better comparison, only values from the depth profile of the reservoir were used. This was not the case for chlorophyll *a* in the Kleine Kinzig Reservoir, as this parameter was only measured in the mixed sample of the reservoir.

### 2.2. Enumeration and identification of bacteria

#### 2.2.1. Microbiological methods

Coliform bacteria and *E. coli* were quantified in 100 ml water sample using the most probable number (MPN) method Colilert^®^-18/Quanti-Tray^®^ (IDEXX Laboratories, Westbrook, USA) according to ISO 9308-2 (2012). For isolation of the coliform bacteria, the wells of the Quanti-Tray^®^ were opened with a sterile scalpel. The liquid was transferred on heterotrophic plate count agar plates (Merck KGaA, Darmstadt, Germany), recommended by German regulations (Deutsches Einheitsverfahren, DEV), with a sterile inoculation loop in order to gain a pure culture. Agar plates were incubated overnight at 36 °C. For identification 10 to 30 isolates per sample were isolated and analyzed with MALDI-TOF MS.

The culture-based detection of enterococci in 100 ml water sample was carried out according to ISO 7899-2 (2000). Additionally, heterotrophic plate counts (HPC) were conducted in 1 ml water sample at 22 °C and 36 °C as described in the German drinking water directive (Drinking Water Ordinance, 2001).

To measure total cell count (TCC), together with high (HNA) and low (LNA) nucleic acid content bacteria, flow cytometry was used (Hammes et al., 2008; Prest et al., 2013). Water samples were diluted in filtered (0.2 µm, Pall Corporation, New York, USA) nuclease-free water (gibco^®^ life technologies^™^, Carlsbad, USA), incubated in the dark with SYBR-Green I (Life Technologies, Eugene, USA, 10,000x in DMSO) for 10 minutes at 37 °C and analyzed with Bio-Rad S3 Cell Sorter (Bio-Rad Laboratories Inc., California, USA) and ProSort™ Software using green fluorescence as a trigger.

#### 2.2.2. Matrix-assisted laser desorption ionization-time of flight mass spectrometry (MALDI-TOF MS)

For the identification of bacterial isolates MALDI-TOF MS was used. Bacterial isolates were picked from DEV agar plates and transferred onto a spot of a polished steel target. Protein extraction was performed with 1 µl of 70% formic acid and then air-dried. The spots were covered with 1 µl of a saturated solution of α-cyano-4-hydroxycinnamic acid (Bruker Daltonics GmbH, Bremen, Germany) solved in 50% acetonitrile and 2.5% trifluoroacetic acid and again air-dried. The plate was applied to the Microflex LT (Bruker Daltonics GmbH, Bremen, Germany) instrument and the spectra were measured according to manufacturer’s instructions (Bruker Daltonics GmbH, Bremen, Germany). For identification MALDI Biotyper and flexControl software were used.

Identification of coliform bacteria with MALDI-TOF MS is often very difficult as they are closely related and MALDI database is not designed for environmental bacteria. Therefore, the database was expanded to ensure accurate identification. Strains, previously identified via MLSA, were added to the database according to manufacturer’s instructions.

#### 2.2.3. Multilocus sequence analysis (MLSA)

Selected isolates were identified at the molecular level via MLSA. For DNA extraction bacterial material was transferred into 50 µl sterile water and then heated for 10 min at 95 °C. After cooling for at least 90 min at - 20 °C, DNA was used for further analysis. MLSA of the genes *atpD*, *infB*, *gyrB* and *rpoB* was performed according to Brady et al. (2013). Amplified PCR products were visualized using QIAxcel^®^ Advanced System (Qiagen, Hilden, Germany) and purified by High Pure PCR Product Purification Kit (Roche, Mannheim, Germany) according to manufacturer’s instructions. Purified DNA was sequenced by StarSEQ^®^ GmbH (Mainz, Germany). Phylogenetic analysis was conducted with MEGA7 Version 7.0.26 (Kumar et al., 2016), using Maximum-Likelihood method and bootstrap values based on 1000 replications.

### 2.3. Bacterial community composition and microbial source tracking

#### 2.3.1. Sample pretreatment and DNA extraction

Water samples (n = 31) for MST and analysis of microbial community were concentrated using membrane filtration (0.2 μm Supor^®^-200 membranes, 47 mm diameter, Pall Corporation, New York, USA). To analyze the fecal markers and the microbial community, total DNA was extracted from the membranes using the FastDNA™ SPIN Kit for Soil (MP Biomedicals, Santa Ana, USA) according to manufacturer’s instructions. DNA quality was evaluated using NanoDrop (Implen, München, Germany).

#### 2.3.2. Quantification of microbial source tracking markers (MST)

Microbial source tracking (MST) is a method to test water samples for a potential fecal pollution and their origin. It is based on the fact that specific microorganisms are associated with a specific host (Hagedorn et al., 2011). Quantitative real-time PCR (qPCR) of 16S rRNA *Bacteroidales* was conducted according to Stange et al. (2019) to identify human, ruminant and pig specific fecal contaminations within the water samples (n = 31), as well as to estimate total amount of fecal contamination from a range of mammals (Layton et al., 2006). Amplified PCR-products were verified using QIAxcel^®^ Advanced System (Qiagen, Hilden, Germany). Efficiency of PCR was between 90 and 105% with a R² value from 0.99. Limit of quantification (LOQ) was 10 copies per reaction.

#### 2.3.3. Analysis of microbial community

For microbiome analysis the V3/V4 region of the 16S rRNA gene was amplified and sequenced at Core Facility Microbiome (Freising, Germany) using Illumina MiSeq^®^ Next Generation Sequencing System. Raw sequence data (n = 869,073) were demultiplexed using Perl script, provided with IMNGS (Lagkouvardos et al., 2016). For further analysis the IMNGS pipeline was used, which implements UPARSE algorithm from the USEARCH8 package (Edgar, 2010, 2013). Sequences were clustered *de novo* to operational taxonomic units (OTUs) with 97% similarity and a relative abundance of ≥ 0.25%. After filtering and checking for chimeras, 504,886 sequences were used for further analysis. Taxonomic classification was carried out using SINA 1.2.11 with a minimum identity query of at least 90% (Pruesse et al., 2012). A Neighbor-Joining tree with 100 bootstraps was conducted using MEGA7 Version 7.0.26 (Kumar et al., 2016). For the analysis of microbial profile the R script Rhea was used (Lagkouvardos et al., 2017).

### 2.4. Statistical Analysis

All statistical analysis were conducted with R (Ihaka and Gentleman, 1996; R Core Team, 2019). Graphics were computed with the packages *ggplot2* (Wickham, 2016) and *ggpubr* (Kassambara, 2020). To determine statistical significant differences Mann-Whitney-U-test was performed.

For plotting the physicochemical parameters bilinear interpolation was conducted using the *akima* package (Akima and Gebhardt, 2016). The relation between physicochemical parameters and coliform bacteria were revealed by the redundancy analysis (RDA) using *zoo* package (Zeileis and Grothendieck, 2005) for linear interpolation of the data and *vegan* package (Oksanen et al., 2019). *R2* was adjusted using Ezekiel’s formula (Ezekiel, 1930) to measure the unbiased amount of explained variation as described by Borcard et al. (2018). ANOVA like permutation test for RDA was performed using *vegan* package (Oksanen et al., 2019) with 999 permutations.

Alpha diversity was estimated using Shannon and Simpson index, where the latter adds more weight to abundant species. The indices were then transformed to effective number of species (Jost, 2006, 2007). Beta diversity examines differences between microbial profiles. Therefore generalized Unifrac distances were calculated using the package *GUniFrac* (Chen, 2018). For visualization a nonparametric multidimensional scaling (nMDS) plot was conducted using the packages *vegan* and *ade4* (Dray and Dufour, 2007; Oksanen et al., 2019).

## 3. Results

### 3.1. Meteorological data

For this study, the two reservoirs were monitored and regularly sampled for two seasons in 2018 and 2019. The main characteristics of the two drinking water reservoirs sampled in this study are summarized in Table 1. The first year of the monitoring study was characterized by major storms in the first quarter of the year, followed by a dry and hot summer. It was the warmest year in Germany since weather records began in 1881 with an average temperature of 10.5 °C and one of the years with the least precipitation (DWD, 2018). For the Kleine Kinzig Reservoir, a mean air temperature of 9.7 °C was measured for the catchment area (Table 2). This represents an increase of 3.1 °C above the long-term average. Precipitation was 14% lower than the years before, but still amounted to 1454 l/m² in 2018. In Klingenberg mean air temperature was 9.2 °C. Total precipitation amounted to 536 l/m², which is significant lower than at the Kleine Kinzig Reservoir (p < 0.001). The second year, 2019, was again hot and dry. With 690 l/m² the precipitation was slightly higher in 2019 compared to 2018 (536 l/m²) at Klingenberg Reservoir, while the average air temperature (9.3 °C) was very similar to 2018. In the Kleine Kinzig Reservoir, mean air temperature was 9.1 °C. Total precipitation was 1940 l/m², which is in the usual range for this area.

**Table 2:** Meteorological and physicochemical parameters of the two reservoirs Klingenberg and Kleine Kinzig.

| Meteorological data |  |  |  |  | Physicochemical data |  |  |  |  |  |  |  |  |  |  |  |
| --- | --- | --- | --- | --- | --- | --- | --- | --- | --- | --- | --- | --- | --- | --- | --- | --- |
|  | PR | WS | AT | WT | PH | OXY | CON | TUR | SAK | CHL | TOC | NO | SI | TP | MN | FE |
|  | l/m<sup>3</sup> | m/s | °C | °C |  | mg/l | µS/cm | FNU | m<sup>-1</sup> | µg/l | mg/l | mg/l | mg/l | mg/l | mg/l | mg/l |
| Klingenberg Reservoir (R1) |  |  |  |  |  |  |  |  |  |  |  |  |  |  |  |  |
| 2018 | 536.0 | 1.0±0.5 | 9.2±8.2 | 9.2±5.0 | 7.2±0.3 | 10.4±1.9 | 146.0±2.6 | 0.5±0.4 | 6.8±0.9 | 3.5±2.0 | 3.5±0.2 | 13.1±2.5 | 7.0±0.5 | 0.025±0.005 | 0.015±0.023 | 0.010±0.004 |
| Surface | - | - | - | 11.4±6.4 | 7.5±0.2 | 10.7±1.5 | 145.7±2.4 | 0.5±0.2 | 6.6±1.0 | 4.6±1.7 | 3.6±0.3 | 12.8±2.4 | 6.8±0.5 | 0.026±0.007 | 0.009±0.009 | 0.010±0.004 |
| Raw | - | - | - | 8.0±4.1 | 7.0±0.3 | 10.0±2.3 | 146.6±2.8 | 1.0±1.2 | 6.9±1.0 | 2.9±2.3 | 3.4±0.2 | 13.2±2.5 | 7.1±0.5 | 0.025±0.005 | 0.029±0.049 | 0.012±0.004 |
| 2019 | 690.4 | 1.0±0.5 | 9.3±7.4 | 8.9±4.8 | 7.1±0.3 | 10.9±2.2 | 157.8±7.9 | 1.0±0.4 | 6.7±0.5 | 7.1±5.4 | 3.5±0.2 | 14.5±3.0 | 5.9±0.5 | 0.021±0.004 | 0.022±0.038 | 0.013±0.006 |
| Surface | - | - | - | 11.6±6.2 | 7.4±0.2 | 11.5±1.9 | 154.9±6.5 | 1.0±0.5 | 6.6±0.6 | 8.6±5.8 | 3.6±0.3 | 13.9±2.8 | 5.6±0.5 | 0.021±0.004 | 0.011±0.007 | 0.012±0.007 |
| Raw | - | - | - | 7.5±3.5 | 6.9±0.3 | 10.0±2.6 | 161.2±7.7 | 1.2±0.6 | 6.8±0.5 | 6.2±5.0 | 3.5±0.2 | 15.0±3.1 | 6.2±0.4 | 0.023±0.004 | 0.060±0.107 | 0.015±0.007 |
| Kleine Kinzig Reservoir (R2) |  |  |  |  |  |  |  |  |  |  |  |  |  |  |  |  |
| 2018 | 1454.0 | 2.5±0.7 | 9.7±7.8 | 6.6±3.7 | 6.8±0.3 | 9.2±2.7 | 40.6±3.2 | 1.3±2.2 | 8.1±2.7 | 6.7±2.1 | 2.2±0.2 | 1.5±0.5 | 1.4±0.4 | 0.034±0.018 | 0.142±0.262 | 0.188±0.244 |
| Surface | - | - | - | 10.6±5.8 | 7.2±0.5 | 10.7±1.5 | 41.5±2.9 | 1.2±0.6 | 6.0±3.0 | 6.7±2.1 | 2.1±0.4 | 0.6±0.5 | 0.8±0.3 | - | 0.019±0.016 | 0.041±0.020 |
| Raw | - | - | - | 4.9±0.6 | 6.6±0.1 | 9.2±2.2 | 39.2±1.9 | 1.2±2.3 | 8.6±1.8 | - | 2.2±0.1 | 1.9±0.1 | 1.6±0.1 | 0.024±0.003 | 0.084±0.079 | 0.136±0.045 |
| 2019 | 1940.6 | 2.4±0.7 | 9.2±7.1 | 6.7±3.2 | 6.7±0.3 | 8.6±1.6 | 39.8±4.2 | 0.6±0.4 | 7.9±1.8 | 5.0±4.8 | 2.4±0.2 | 1.5±0.4 | 1.5±0.1 | 0.017±0.004 | 0.026±0.041 | 0.046±0.030 |
| Surface | - | - | - | 9.5±5.2 | 6.9±0.3 | 9.7±1.2 | 39.8±4.4 | 0.6±0.3 | 7.0±2.6 | 6.4±3.9 | 2.2±0.2 | 1.1±0.6 | 1.4±0.2 | - | 0.011±0.009 | 0.027±0.016 |
| Raw | - | - | - | 5.4±1.2 | 6.7±0.2 | 8.4±1.4 | 39.6±4.3 | 0.5±0.4 | 8.4±1.3 | - | 2.5±0.1 | 1.7±0.1 | 1.5±0.1 | 0.017±0.002 | 0.020±0.019 | 0.042±0.009 |
| Differences between the two reservoirs (Mann-Whitney-U-test) |  |  |  |  |  |  |  |  |  |  |  |  |  |  |  |  |
| Surface | *** | *** | NS | NS | *** | *** | *** | NS | NS | NS | *** | *** | *** | - | ** | *** |
| Raw | *** | *** | NS | *** | *** | *** | *** | *** | *** | - | *** | *** | *** | ** | *** | *** |

Values show means and standard derivation, except for precipitation that shows sum of daily precipitation. PR = precipitation, WS = wind speed, AT = air temperature, WT = water temperature, pH = pH-value, OXY = oxygen, CON = conductivity at 25 °C, TUR = turbidity, SAK = SAK 254 nm, CHL = chlorophyll a, TOC = total organic carbon, NO = nitrate, SI = silicate, TP = total phosphate, MN = total manganese, FE = dissolved iron. Statistical significance is represented by \* (p < 0.05), \*\* (p < 0.01) and \*\*\* (p < 0.001). NS means no significance.

### 3.2. Physicochemical parameters

During the whole investigation period, physicochemical parameters were analyzed. An overview is given in Table 2 for both reservoirs. Selected parameters (temperature, pH, O2 content) are shown in Fig. 1.

**Fig. 1:**
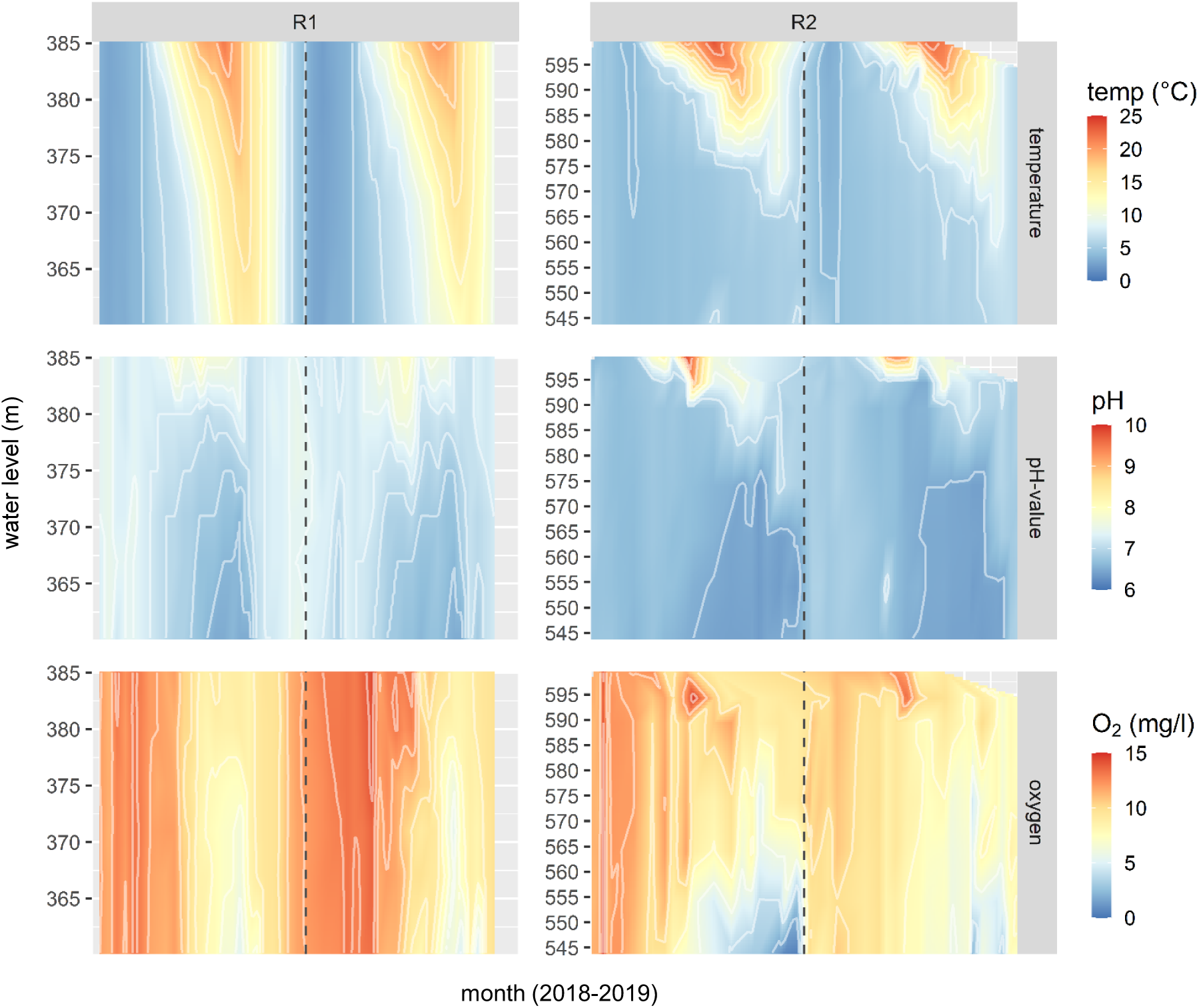
Physical-chemical parameters in the Klingenberg (R1) and Kleine Kinzig Reservoir (R2) in 2018 and 2019. The graph shows the measured values for temperature, pH and O2 in the depth profile of the two reservoirs over time.

The two reservoirs were selected since they are comparable in size, but differ in other parameters and represent a typical mesotrophic (Klingenberg) and oligotrophic (Kleine Kinzig) drinking water reservoir in Germany. Significant differences can be observed regarding their metal and nutrient content (Table 2). For example, TOC, nitrate as well as silicate content is higher in Klingenberg Reservoir (p < 0.001), whereas total manganese and dissolved iron content is higher in Kleine Kinzig Reservoir (p < 0.01). Furthermore, Kleine Kinzig Reservoir as a humic material shaped reservoir has a higher SAK (254 nm).

The two reservoirs did not only differ with regard to the nutrient level, but also in water temperature. Klingenberg Reservoir is noticeably warmer with a mean water temperature 2 to 3 °C higher than in Kleine Kinzig Reservoir (Table 2), mainly due to higher temperatures in the deeper water layers (Fig. 1) (p < 0.001).

Stratification duration is another difference between the two reservoirs, caused by the differences in water temperature. For both reservoirs and years, stratification started in April, but it stopped much earlier in Klingenberg with the beginning of October, when temperature uniformity was reached and all layers had high water temperature. In Kleine Kinzig Reservoir, the stratification period lasted longer, until November in 2018 and until January 2020 in the 2019 season.

### 3.3. Seasonal changes of microbiological parameters

Regular microbiological monitoring of surface water and raw water (used for drinking water production) included the parameters *E. coli,* coliform bacteria, enterococci, heterotrophic plate counts (at 22 °C and 36 °C) and total cell counts. Fig. 2 gives an overview about the measured numbers of these parameters, median values of coliform bacteria, enterococci and *E. coli* are shown in the Supplement (Table S1).

**Fig. 2:**
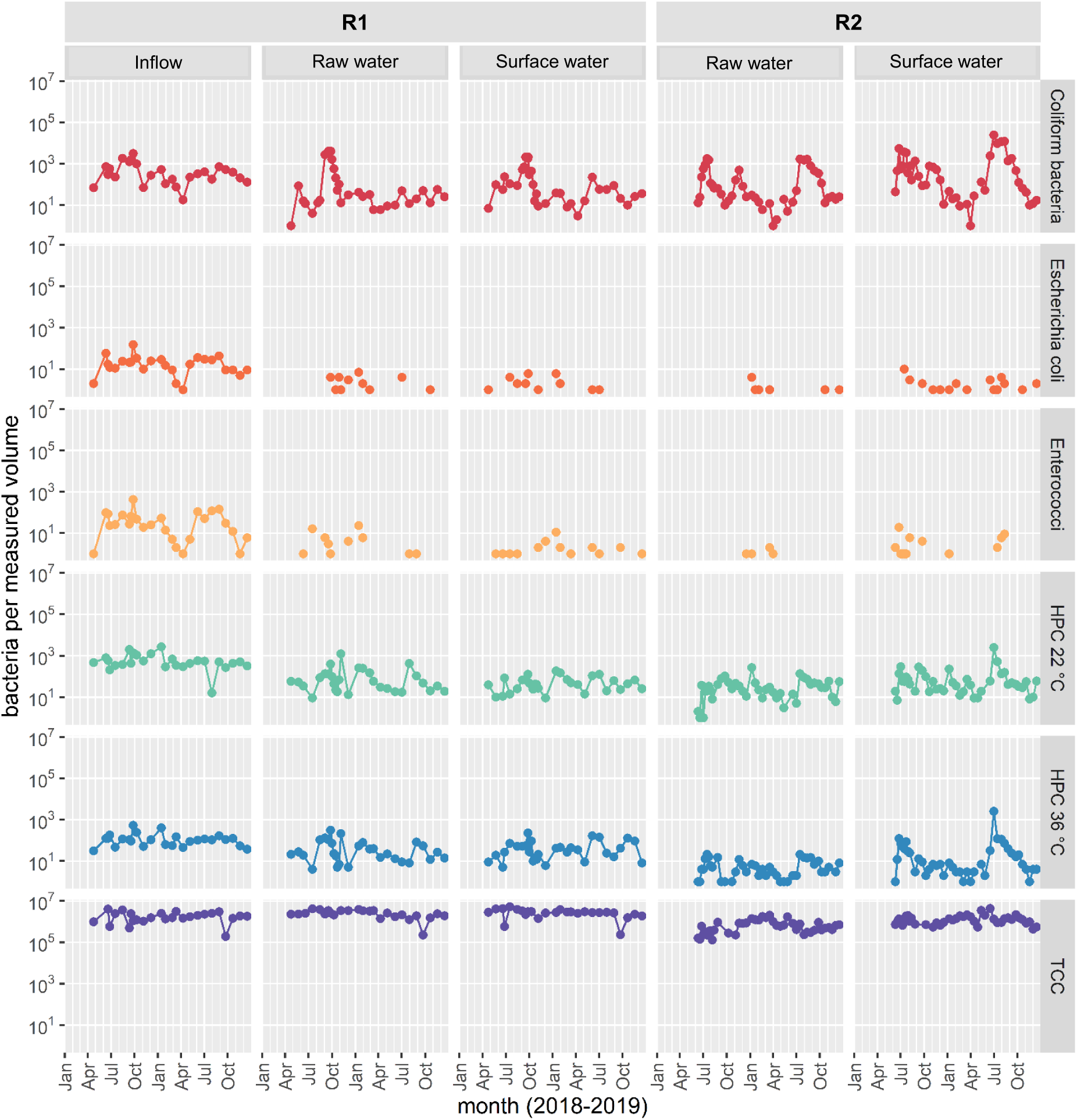
Microbiological parameters in Klingenberg (R1) and Kleine Kinzig (R2) reservoirs in 2018 and 2019. The investigated volume was 100 ml for coliform bacteria, *Escherichia coli* and enterococci and 1 ml for HPC at 22 °C and 36 °C and the total cell count (TCC).

In Kleine Kinzig Reservoir, 44 samples of each water were analyzed during the monitoring study. In the majority of the samples, the FIB *E. coli* and enterococci could not be detected in 100 ml water samples. With an abundance of 34% respectively 27% of the samples they were detected more often in the surface water samples than in the raw water samples (14% and 9%, respectively). Maximum numbers of *E. coli* were 10 MPN/100 ml in the surface water and 4 MPN/100 ml in raw water. For enterococci, the highest levels detected were 19 CFU/100 ml in the surface water and 2 CFU/100 ml in the raw water. Dominating species was the environmental enterococcus strain *Enterococcus rotai* (61% of positive results).

In contrast, coliform bacteria were detected in all except two samples (98%). As shown in Fig. 2, the concentration of coliform bacteria varied over four orders of magnitude ranging from below detection limit in 100 ml to above 2.4 x 10^4^ MPN/100 ml. Periods of high concentrations have been observed in both years 2018 and 2019, starting from the end of June until the end of September. While the concentration of coliform bacteria remained high in 2019 over the entire summer period, a decrease in number could be observed in 2018 after the initial peak in June/July and a second peak occurred in November 2018. Highest numbers in coliform bacteria were observed in the surface water with a maximum of 5.4 x 10^3^ MPN/100 ml in June 2018 and about ten times the amount in July 2019 (>2.4 x 10^4^ MPN/100 ml). In the raw water, numbers were lower with a maximum number of about 1.7 x 10^3^ MPN/100 ml in both years.

In Klingenberg Reservoir regular sampling was conducted 32 times. The fecal indicators *E. coli* and enterococci could be detected more often than in the other reservoir. They were detected in 31% respectively 59% of all surface water samples and in 34% respectively 50% of all raw water samples. The highest values of *E. coli* were 7 MPN/100 ml, both in surface and raw water. Enterococci reached maximum numbers of 11 CFU/100 ml in raw water and 23 CFU/100 ml in surface water.

Coliform bacteria were detected in all samples. In contrast to Kleine Kinzig Reservoir, the highest numbers of coliform bacteria were reached in the raw water with 4.0 x 10^3^ MPN/100 ml in September 2018. In the surface water, the maximum numbers were also detected in September 2018, but reached with 2.1 x 10^3^ MPN/100 ml a lower level. Surprisingly, these high numbers of coliforms were only observed in 2018. In 2019, the maximum number of coliform bacteria was 2.3 x 10^2^ MPN/100 ml, reached in September in the surface water.

In Klingenberg, the main inflow into the reservoir (Wilde Weißeritz) was also sampled regularly (n = 26). In contrast to the reservoir (surface and raw water), the inflow did not show high seasonal differences. Other than in the reservoir, FIB *E. coli* and enterococci were detected in all samples. They reached maximum values of 1.5 x 10^2^ MPN/100 ml for *E. coli* and 4.2 x 10^2^ CFU/100 ml for enterococci, thus around 1.5 orders of magnitude higher than in the reservoir. Coliform bacteria were observed in all samples, too. Here the maximum number was 3.1 x 10^3^ MPN/100 ml (Fig. 2).

### 3.4. Identification of coliform bacteria

Seasonal dynamics were not only found for the number of coliform bacteria, but also for the species diversity, as shown in Fig. 3. Overall, 45 species of coliform bacteria, belonging to 20 genera, could be identified. The five most abundant genera were *Serratia, Enterobacter, Lelliottia, Citrobacter* and *Buttiauxella*.

**Fig. 3:**
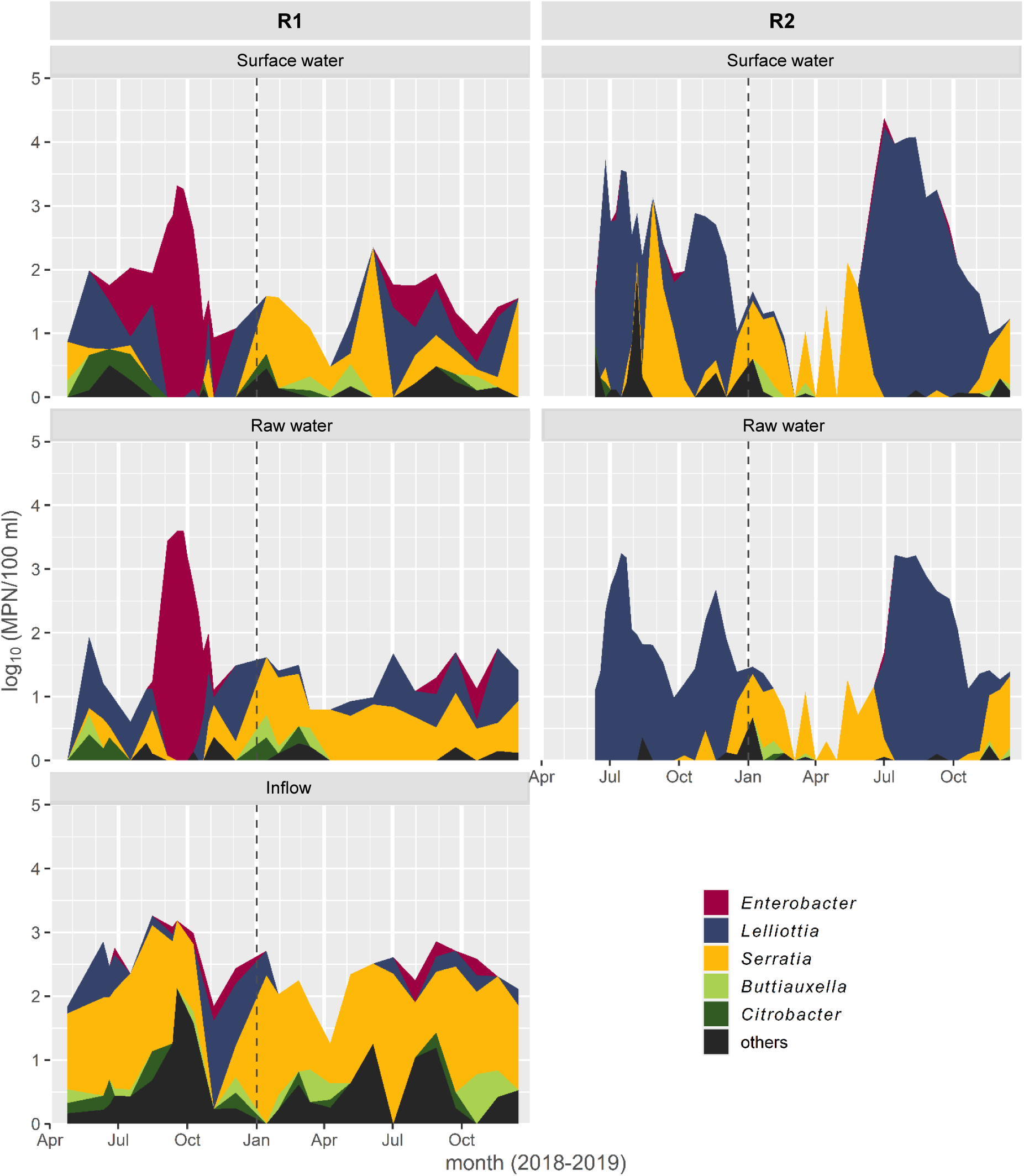
Number and genera of the coliform bacteria in the Klingenberg (R1) and Kleine Kinzig Reservoir (R2) in 2018 and 2019. “Others” include the genera: *Dickeya, Hafnia, Klebsiella, Kluyvera, Kosakonia, Leclercia, Moellerella, Morganella, Pantoea, Plesiomonas, Proteus, Providencia, Rahnella, Raoultella* and *Yersinia*.

In Kleine Kinzig Reservoir at least 26 species of 11 genera could be identified. Mostly two genera of coliform bacteria dominated, namely *Lelliottia* (72%) and *Serratia* (21%). During winter, mainly *Serratia* occurred, but as soon as concentrations of coliform bacteria increased, *Lelliottia* was dominant. By measuring the species diversity using Simpson effective (Table S2), the diversity decreased during mass proliferation from three to four effective species to only two, namely *Lelliottia amnigena* and *Lelliottia aquatilis*.

The Klingenberg Reservoir differed considerably. In the reservoir 28 species out of 12 genera could be observed. As in Kleine Kinzig Reservoir *Serratia* dominated during winter period, *Lelliottia* also occurred frequently, but the genus did not dominate. Instead, *Enterobacter asburiae* dominated here during late summer 2018, the time with the highest concentrations of coliform bacteria. Species diversity showed an even higher change during this time as it decreased from about six effective species to only one. In this monitoring study, the first occurrence of *Enterobacter asburiae* was in June/July of both years in the surface water. In August, *Enterobacter asburiae* was observed in the raw water. While this species could only be identified sporadically in 2019, it dominated in 2018 from September onwards throughout the whole reservoir.

In contrast, the main inflow did not change much over the sampling period. Richness was much higher here, with 43 species out of 18 genera. The dominating genus was *Serratia*. In cases where *Enterobacter* was detected, it was mainly identified as *Enterobacter cloacae*. In the inflow, no change in species diversity could be observed during times with high concentrations of coliform bacteria in the reservoir compared to times with low concentrations. The diversity was about six to seven effective species all over the years, thus higher than in the reservoir.

### 3.5. Sampling campaign

At Klingenberg Reservoir, two sampling campaigns were conducted in addition to the monitoring program in 2018, one in June and the other one in September. At that time, not only the six depth layers and the main inflow were sampled, but also the whole water body of the reservoir together with additional inflows. Fig. 4 gives an overview about the results of the quantification and identification of coliform bacteria for exemplary presented samples.

**Fig. 4:**
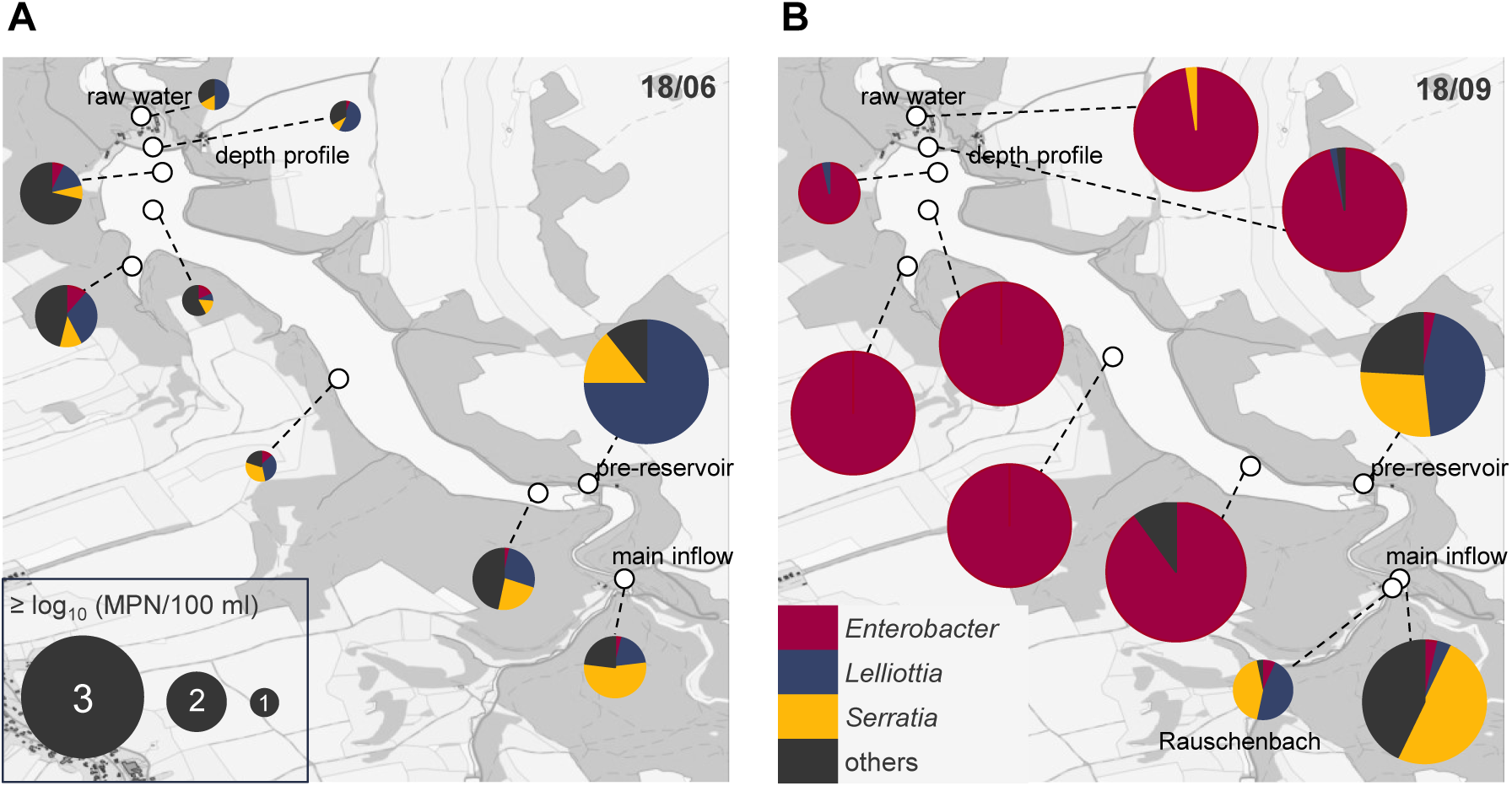
Quantification and identification of coliform bacteria at the Klingenberg Reservoir during sampling campaign in June (A) and September 2018 (B).

In June 2018 in addition to the six regular samples, 13 samples from the surface of the reservoir were sampled (n = 19). Furthermore, together with the main inflow (Wilde Weißeritz), another small inflow and the outflow of the pre-reservoir could be sampled (n = 3). All other tributaries had low water levels due to the drought during this year. As presented in Fig. 4A in the water of the reservoir, numbers of coliform bacteria were generally low with a median of 33 MPN/100 ml (n = 19). Only in the inflow higher numbers of coliform bacteria were measured, e.g. 5.8 x 10^2^ MPN/100 ml at the main inflow, and 2.6 x 10^3^ MPN/100 ml at the outflow of the pre-reservoir.

From selected samples, coliform bacteria were identified (n = 11). Various genera were present in the reservoir water, like *Lelliottia* spp. (23%), *Serratia* spp. (19%), *Citrobacter* spp. (14%) and *Enterobacter* spp. (11%), most of them being identified as *Enterobacter asburiae*. In the main inflow, *Serratia* spp. dominated (56%), whereas in the pre-reservoir *Lelliottia* spp. was dominant (75%). The genus *Enterobacter* was only present in low abundance (4%) in the main inflow with all isolates identified as *Enterobacter cloacae*.

In September 2018, 14 additional samples from the water body of the reservoir were sampled, together with the depth profile (n = 20). Furthermore, the main inflow, the outflow of the pre-reservoir, as well as an additional water pipe from the Rauschenbach Reservoir were sampled (n = 3). During that time, additional water was transferred from the Rauschenbach Reservoir into the Klingenberg Reservoir in order to prevent the reservoir from carrying too little water. Results are presented in Fig. 4B.

The observed numbers of coliform bacteria in the reservoir water (n = 20) were much higher than in June (median: 1.4 x 10^3^ MPN/100 ml), nearly all of them being identified as *Enterobacter asburiae* (95%). In the inflowing water, numbers of coliform bacteria reached 7 x 10^2^ to 1.6 x 10^3^ MPN/100 ml. Similar to June, *Serratia* spp. dominated in the main inflow (52%) and *Lelliottia* spp. in the pre-reservoir (43%). In the pipe from Rauschenbach Reservoir, both genera had approximately the same frequencies (>40%). Less than 5% of all isolates from the inflows were identified as *Enterobacter* spp. (mainly *Enterobacter ludwigii*).

### 3.6. Microbial Source Tracking (MST)

The detection of MST markers indicates the potential source of hygienically-relevant microbial load in the reservoir. Therefore, samples were taken at two time points in each reservoir, once with high numbers of coliform bacteria, once without. For Klingenberg Reservoir in June and September 2018 surface water of the complete reservoir, the depth profile and all inflows were sampled, as part of the sampling campaign. In Kleine Kinzig Reservoir, sampling was conducted in July and December 2019 from the depth profile. With extracted DNA of the samples, qPCR with host-specific 16S rRNA sequences of *Bacteroidales* bacteria was carried out, to identify human-, ruminant- and pig-specific fecal contamination within the samples. These markers were compared to the total cell counts obtained by flow cytometry measurements as well as the total 16S rRNA genes quantified by qPCR (Table 3).

**Table 3:** Mean values and standard deviation of total cell count (TCC), and gene copies of 168 rRNA genes and microbial source tracking markers genes (in 1 mL)

|  |  | **Klingenberg Reservoir (R1)** |  | **Kleine Kinzig Reservoir (R2)** |  |
| --- | --- | --- | --- | --- | --- |
|  |  | **low** | **high** | **low** | **high** |
| **number of samples** |  | 15 | 5 | 5 | 6 |
| **date** |  | June 2018 | September 2018 | December 2019 | July 2019 |
| **TCC** | counts/ml | $7.4 \times 10^5 \pm 6.1 \times 10^4$ | $8.2 \times 10^5 \pm 1.2 \times 10^5$ | $5.4 \times 10^5 \pm 1.1 \times 10^5$ | $5.7 \times 10^5 \pm 3.4 \times 10^5$ |
| **LNA** | % | 37.2% | 31.8% | 15.5% | 12.9% |
| **16S rRNA** | GC/ml | $1.3 \times 10^6 \pm 9.8 \times 10^4$ | $1.3 \times 10^6 \pm 1.5 \times 10^5$ | $6.3 \times 10^5 \pm 5.3 \times 10^4$ | $5.9 \times 10^5 \pm 1.7 \times 10^5$ |
| **unspecific** | GC/ml | $1.1 \times 10^2 \pm 5.2 \times 10^1$ | $2.0 \times 10^3 \pm 6.5 \times 10^2$ | $2.8 \times 10^3 \pm 3.6 \times 10^2$ | $3.8 \times 10^3 \pm 1.5 \times 10^3$ |
| **human-specific** | GC/ml | <LOQ | <LOQ | <LOQ | <LOQ |
| **ruminant-specific** | GC/ml | <LOQ | <LOQ | <LOQ | <LOQ |
| **pig-specific** | GC/ml | <LOQ | $4.0 \times 10^2$ * | <LOQ | <LOQ |

\* only measured in one sample, therefore no standard error

Altogether, the total amount of bacteria within the samples was around 10^6^ bacteria per ml. TCC for all measured samples (n = 24) had a mean concentration of 7.3 x 10^5^ bacteria per ml, the measured 16S rRNA genes over all samples (n = 31) a mean concentration of 1.1 x 10^6^ genes/ml (Table 3).

Unspecific marker genes, to measure all *Bacteroides* in the sample and therefore total amount of fecal contamination, made around 0.2% of all 16S rRNA (2 x 10³ gene copies/ml). Neither human-nor ruminant-specific fecal marker genes were detected in any of the samples (<LOQ). Out of the total number of 31 samples, pig-specific fecal markers could only be detected once, in the main inflow (Wilde Weißeritz) of Klingenberg Reservoir in September with a concentration of 4 x 10² genes/ml (Table 3).

### 3.7. Microbial community

To compare the microbial community between the two reservoirs, as well as between times with high and low concentrations of coliform bacteria, 16S rRNA gene amplicon sequencing was conducted. Therefore water samples were taken at two time points: with high and with low concentrations of coliform bacteria in the two reservoirs. For Klingenberg Reservoir this was June (low) and September 2018 (high), for Kleine Kinzig July (high) and December 2019 (low).

Overall 869,073 sequences from 31 water samples were demultiplexed. After checking for quality and chimeras 504,886 sequences were further analyzed. Altogether, 362 OTUs were observed within the water samples. Alpha diversity showed a richness of 225 ± 6 OTUs per sample (Table S3). According to Shannon 72 ± 4 effective OTUs per sample occurred. Adding more weight to abundant species, Simpson measures 39 ± 3 effective OTUs per sample. Both, richness and diversity was slightly higher in Kleine Kinzig Reservoir than in Klingenberg Reservoir.

The OTUs could be assigned to 17 phyla (Fig. 5A). The three most abundant ones were *Actinobacteriota* (32.9%), *Proteobacteria* (31.2%) and *Bacteroidota* (20.4%). Fig. 5B gives an overview about the order assigned to these three phyla. The most abundant order was *Frankiales* (*Actinobacteraeota*) with the family *Sporichthyaceae* (25.9%). The most abundant order within *Proteobacteria* was *Burkholderiales* with the family *Comamonadaceae*. The order to which the coliform bacteria belong (*Enterobacterales*) was not very common with only 0.04%. By comparing the two reservoirs with Kruskal-Wallis Rank Sum Test it could be observed that *Bacteroidota* were significantly more abundant in Klingenberg than in Kleine Kinzig Reservoir (*p* < 0.001), whereas the *Planctomycetota* were significantly more abundant in the latter (*p* < 0.001) (data not shown).

**Fig. 5:**
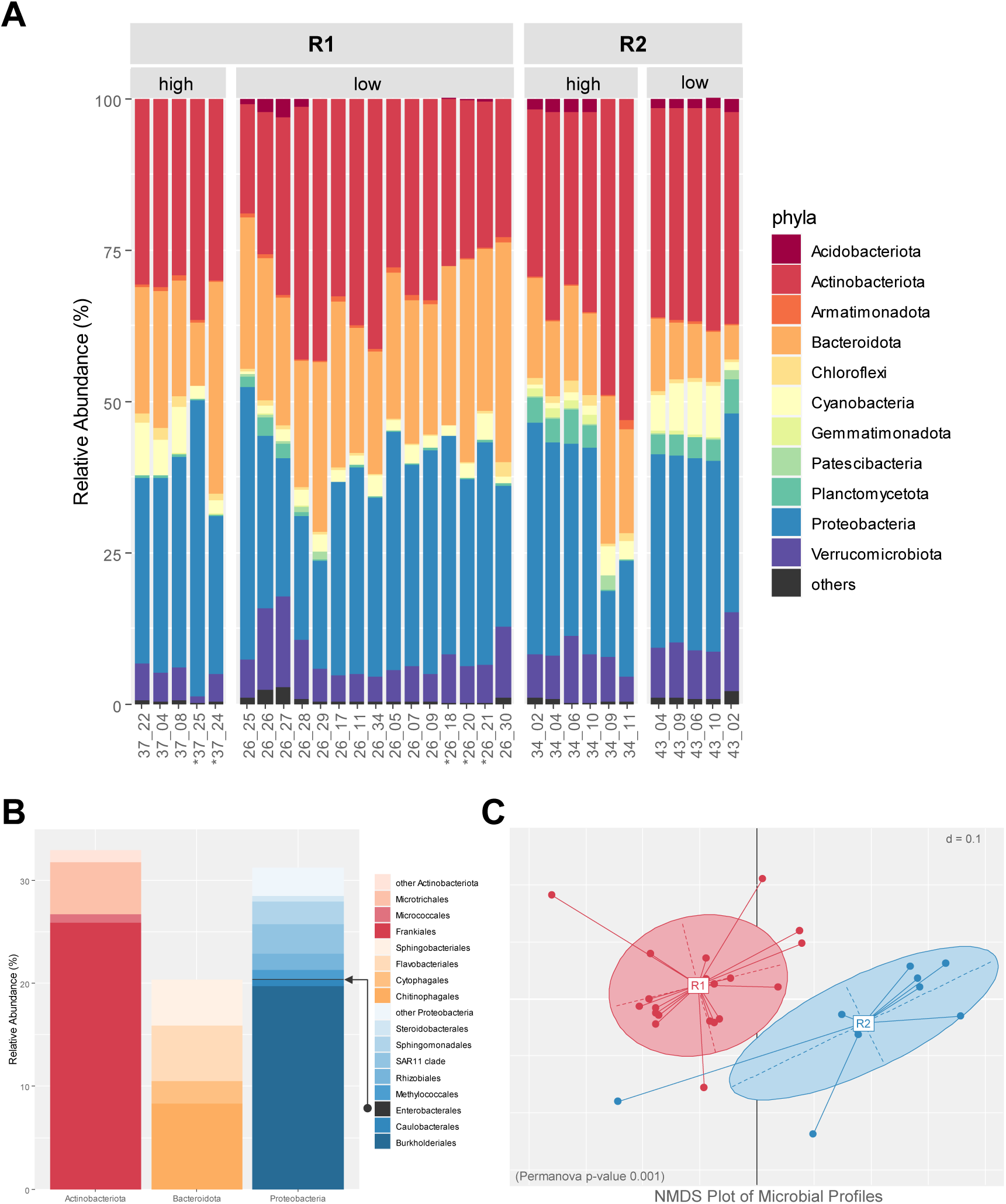
Analysis of the microbial community in the Klingenberg (R1) and Kleine Kinzig Reservoir (R2). Samples marked with * represent inflows (A). Detailed presentation of the three most abundant phyla *Actinobacteriota*, *Bacteroidota* and *Proteobacteria* (B). The *Enterobacterales* are highlighted with the black arrow. Two-dimensional nonparametric multidimensional scaling (NMDS) of the two reservoirs based on generalized UniFrac distances (C).

Nonparametric multidimensional analyses of phylogenetic distances between the samples revealed that clusters were built according to the reservoir (Fig. 5C). Nevertheless, no significant differences could be observed between time points with high or with low numbers of coliform bacteria.

### 3.8. Redundancy Analysis (RDA)

To define parameters affecting the proliferation of coliform bacteria, RDA was conducted using data from surface and raw water (Fig. 6). Permutation test for RDA revealed different factors explaining the variance of coliform species in Klingenberg (Fig. 6A) as well as Kleine Kinzig Reservoir (Fig. 6B), e.g. environmental factors (water temperature, oxygen, pH value and SAK) as well as metals and nutrients (manganese, iron, TOC) and microbiological parameters (TCC). Furthermore, some factors were only significant for the variation in one of the two reservoirs: air temperature, chlorophyll, total phosphate, silicate, nitrate, HPC at 22 °C (Klingenberg) and conductivity and turbidity (Kleine Kinzig). For both reservoirs, depth was no significant factor for species composition.

**Fig. 6:**
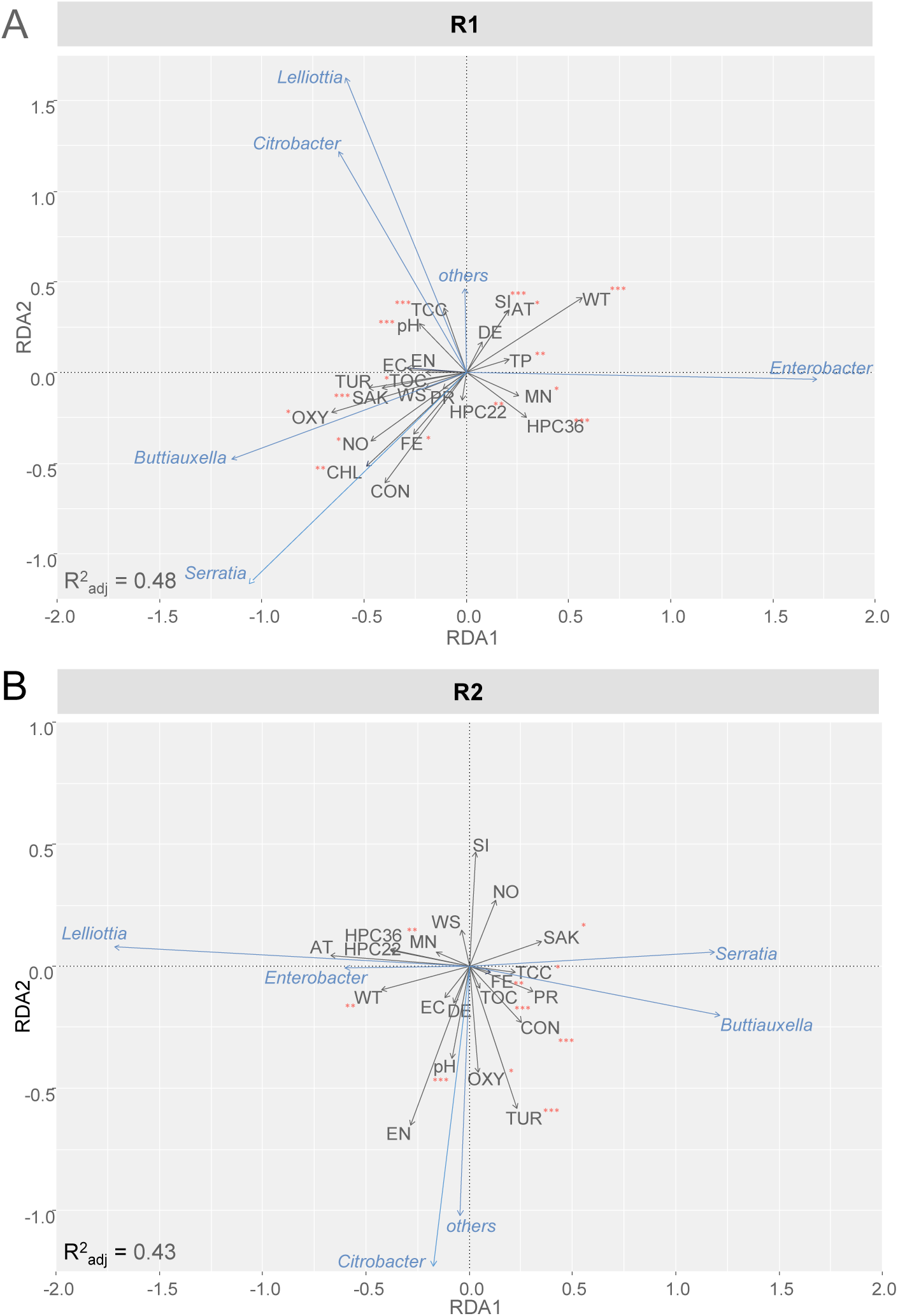
Redundancy Analysis to reveal relationship between physicochemical parameters and coliform bacteria in Klingenberg Reservoir (A) and Kleine Kinzig Reservoir (B). DE = depth, PR = precipitation, WS = wind speed, AT = air temperature, WT = water temperature, pH = pH-value, OXY = oxygen, CON = conductivity at 25 °C, TUR = turbidity, SAK = SAK 254 nm, CHL = chlorophyll a, TOC = total organic carbon, NO = nitrate, SI = silicate, TP = total phosphate, MN = manganese, FE = iron. Statistical significance is represented by * (p < 0.05), ** (p < 0.01) and *** (p < 0.001).

Factors correlating positively with the proliferating coliform species were in both reservoirs the water temperature as well as manganese content. Both were additionally negatively correlated with SAK and TOC. Furthermore, *Enterobacter* spp. in Klingenberg Reservoir were correlated with total phosphate (positive) and oxygen content (negative). The occurrence of *Lelliottia* spp. in Kleine Kinzig Reservoir was negatively correlated with *Serratia* spp., iron, as well as TCC.

### 3.9. Other reservoirs

In this study, the two genera *Enterobacter* and *Lelliottia* were identified as being present in drinking water reservoirs during times with high concentrations of coliform bacteria, thus, assumingly being capable to proliferate within these reservoirs, depending on the reservoir being investigated (see discussion below). Therefore, the question arose how the situation was in other drinking water reservoirs and lakes, where mass proliferation of coliform bacteria has been observed.

Water samples and isolates of coliform bacteria from different lakes and reservoirs in Germany were analyzed. The results are summarized in Table 4. The phylogenetic analysis of the isolated strains by MLSA is shown in Fig. 7. It turned out that *Enterobacter asburiae* could be detected in several other reservoirs and lakes during times with high numbers of coliform bacteria, like in Lake Constance, Rappbode Reservoir, located in the Harz Mountains in Saxony-Anhalt, and Breitenbach Reservoir in North Rhine-Westphalia. *Lelliottia amnigena* could be detected and identified in Söse Reservoir, located in the Harz Mountains in Lower Saxony during mass proliferation.

**Fig. 7:**
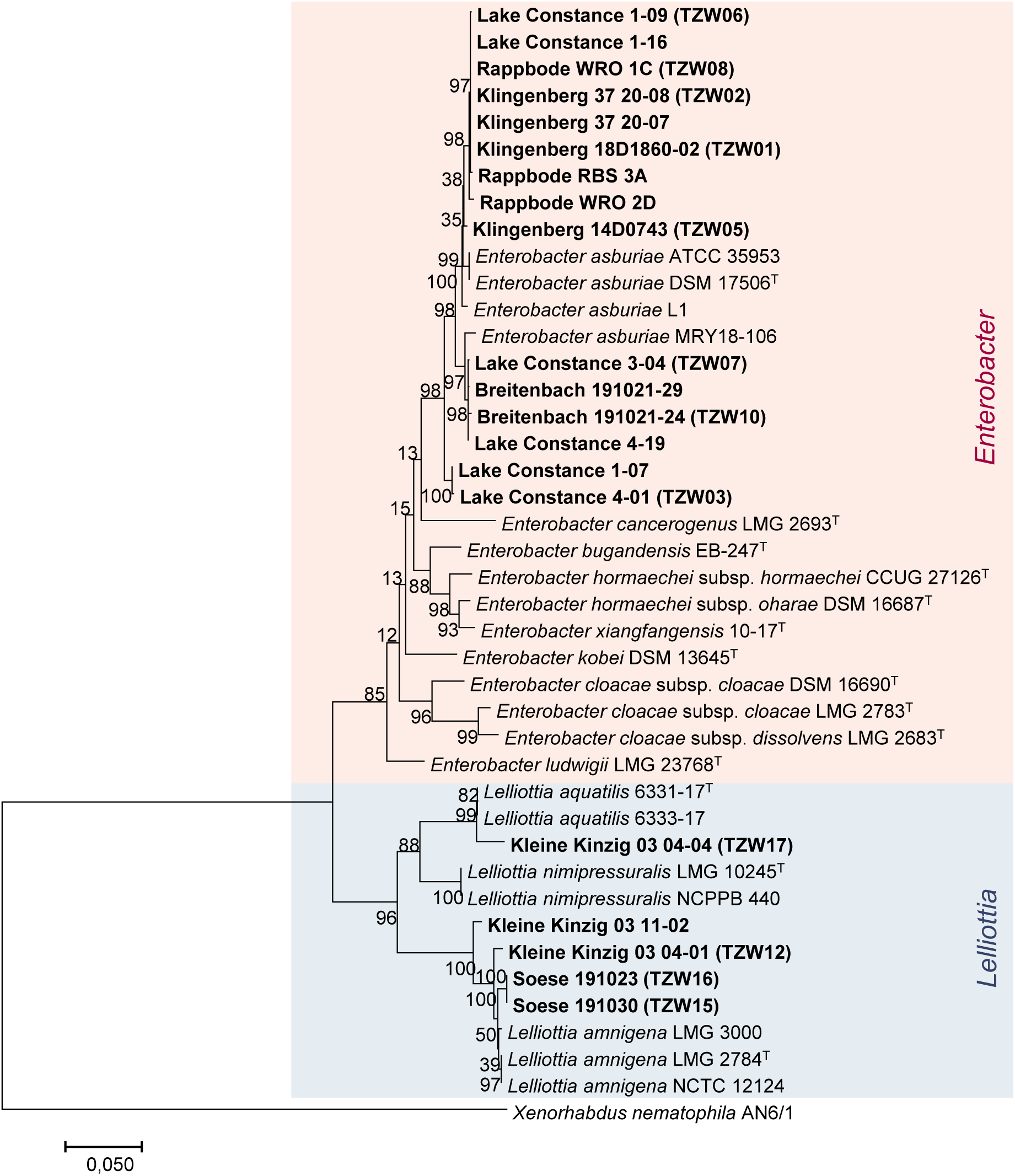
Phylogenetic analysis of *Enterobacter* and *Lelliottia* spp. from different drinking water reservoirs and Lake Constance. Maximum likelihood tree based on the genes *atpD*, *gyrB*, *infB* and *rpoB* (MLSA-PCR) with 1000 bootstraps. As an outgroup *Xenorhabdus nematophila* AN6/1 was used. The reference bar represents a difference of the sequences of 5%.

**Table 4:** Description of additional drinking water reservoirs and lakes with mass proliferation of coliform bacteria studied.

|  | Lake Constance | Rappbode Reservoir | Breitenbach Reservoir | Söse Reservoir |
| --- | --- | --- | --- | --- |
| **Description of surface water** |  |  |  |  |
| **State** | Germany, Austria, Switzerland | Saxony-Anhalt | North Rhine-Westphalia | Lower Saxony |
| **Location** | Foothills of the Alps | Harz Mountains | Rothaargebirge | Harz Mountains |
| **River basin** | Rhine | Elbe | Rhine | Weser |
| **Altitude** | 395 m ASL | 423 m ASL | 370 m ASL | 326 m ASL |
| **Depth (Average)** | 90 m | 29 m | 14 m | 21 m |
| **Depth (Maximum)** | 251 m | 89 m | 35 m | 49 m |
| **Water surface** | 53,000 ha | 390 ha | 58 ha | 124 ha |
| **Capacity** | 48 Mrd m<sup>3</sup> | 110 Mio m<sup>3</sup> | 8 Mio m<sup>3</sup> | 25 Mio m<sup>3</sup> |
| **Raw water delivery** | 5500 l/s | 2900 l/s | 190 l/s | 550 l/s |
| **Catchment area** | 11,500 km<sup>2</sup> | 115 km<sup>2</sup><br>(mainly forest) | 12 km<sup>2</sup><br>(mainly forest) | 49 km<sup>2</sup> |
| **Ecology** | oligotrophic | mesotrophic | oligo-/mesotrophic | oligotrophic |
| **Geology** | moraines | granite | shale, graywacke | shale, granite |
| **Proliferation of coliform bacteria** |  |  |  |  |
| **Method** | Colilert | Colilert | Colilert | CCA |
| **Concentration (Maximum)** | >200 MPN/100 ml (R, S) | >2420 MPN/100 ml (S) | 7940 MPN/100 ml (R) | >200 CFU/100 ml (R) |
| **Date (Maximum)** | 27.08./16.09.2015 (R, S) | 01./09.08.2018 (S) | 18.09.2019 (R) | 22.07.2019 (R) |
| **Concentration (Sampling)** | 172 MPN/100 ml (R) | >2420 MPN/100 ml (S),<br>1986 MPN/100 ml (R) | 326 MPN/100 ml (R) | 264 CFU/100 ml (R) |
| **Date (Sampling)** | 02.11.2015 (R) | 09.08.2018 (S),<br>28.08.2018 (R) | 21.10.2019 (R) | 30.10.2019 (R) |
| **Identification (Sampling)** | *Enterobacter asburiae* | *Enterobacter asburiae* | *Enterobacter asburiae* | *Lelliottia amnigena* |

S: Surface water; R: Raw water

## 4. Discussion

### 4.1. Mass proliferation of coliform bacteria during the summer months is dominated by *Enterobacter asburiae* and *Lelliottia* spp.

In this study, seasonal dynamics of coliform bacteria in two reservoirs in Germany were investigated. The two reservoirs were selected, as high numbers of coliform bacteria have been observed in previous years. Both reservoirs are similar in size, but apart from that are different and can be regarded as representative for typical mesotrophic (Klingenberg) and oligotrophic (Kleine Kinzig) drinking water reservoirs.

During summer months, numbers of coliform bacteria suddenly increased in both reservoirs and reached densities of 10^3^ MPN/100 ml and more in the whole depth profile of the reservoir as well as the entire water body. Not only the number but also the species composition changed entirely. Diversity decreased to only one or two species of coliform bacteria. Therefore, we assume that this sudden increase of single or a few species can be considered as a mass proliferation of coliform bacteria or “coliform bloom” within the reservoir (see also further discussion below).

Depending on the reservoir *E. asburiae, L. amnigena* or *L. aquatilis,* respectively, were dominating. Interestingly, closely related strains were found in different reservoirs all over Germany (Fig. 7). They are well known to occur in water, e.g. *L. amnigena* (formerly *Enterobacter amnigenus*) and *L. aquatilis* were first described as being isolated from water samples (Izard et al., 1981; Kämpfer et al., 2008; Kämpfer et al., 2018). *Enterobacter asburiae* belongs to the *Enterobacter cloacae* complex, a complex consisting of clinical as well as environmental isolates, whose taxonomy remains complicated and differentiation is difficult (Brady et al., 2013; Hoffmann and Roggenkamp, 2003; Mezzatesta et al., 2012). The species was initially isolated from clinical material, however its clinical significance is still uncertain since it also occurs in environmental samples like water (Brenner et al., 1986; Davin-Regli et al., 2019; Kämpfer et al., 2008; Sanders and Sanders, 1997; Suzuki et al., 2018). Furthermore, proliferation of *Enterobacter* spp. has been described in drinking water distribution systems (Edberg et al., 1994).

### 4.2. *Frankiales* and *Burkholderiales* dominate the microbial community

To see effects of proliferation events on the microbial community, 16S rRNA amplicon analysis was conducted. The three most abundant phyla were *Actinobacteriota*, *Bacteroidota* and *Proteobacteria*. All three phyla are well known to occur in drinking water samples as well as freshwater habitats (Paruch et al., 2019; Perrin et al., 2019; Pinto et al., 2014; Rodriguez-R et al., 2020; Tamames et al., 2010; Vaz-Moreira et al., 2014; Zeng et al., 2013). *Firmicutes*, which on the contrary are frequently measured in gastrointestinal microbiome and are reported as one of the major human fecal bacteria in watershed, were not abundant (Paruch et al., 2019; Unno et al., 2010).

*Actinobacteriota* are Gram-positive bacteria distributed in aquatic as well as terrestrial ecosystems and constitute one of the largest bacterial phyla (Barka et al., 2016). They have a mycelial lifestyle and an extensive secondary metabolism. *Frankiales* are known to dominate *Actinobacteriota* in water samples (Pinel et al., 2020). *Frankia* is a nitrogen-fixing actinobacterium and is associated with plants (Barka et al., 2016; Rosenberg et al., 2014). *Actinobacteriota* have previously been described as small bacteria with LNA-content (Proctor et al., 2018). This is consistent with the high numbers of LNA-content bacteria detected during this study amounting to about 75% (Table 3).

*Proteobacteria* have been reported to be the most abundant bacteria in epilimnic waters, especially *Burkolderiales* dominate in less productive reservoirs like oligotrophic and mesotrophic ones (Llirós et al., 2014). They can be associated with cyanobacteria or particles and have been correlated to DOC and nitrate concentration (Llirós et al., 2014). *Burkholderiales* are a diverse order, belonging to the β-*Proteobacteria*, including strictly aerobic and facultative anaerobic chemoorganotrophs, obligate and facultative chemolithothrophs, nitrogen-fixing organisms, as well as plant, animal and human pathogens (Garrity et al., 2005). The family most abundant during this study was the chemoorganotrophic or facultative chemolithothrophic *Comamonadaceae,* where isolates have been obtained from various sources including soil, water and environment (Garrity et al., 2005; Pinto et al., 2014; Tamames et al., 2010).

Within the *Proteobacteria* it was expected to detect the order *Enterobacterales* frequently, especially during times with high concentrations of coliform bacteria. Nevertheless, this was not the case. They only had a frequency of 0.04%. By comparing the maximum concentration of coliform bacteria (2.4 x 10^4^ MPN/100 ml) with the TCC and 16S rRNA genes, even these high concentrations of coliform bacteria make only around 0.02% of the total bacterial cells. Therefore it is not surprising, that no significant differences of the microbial community could be found between samples with high and low coliform bacteria numbers.

Instead, significant differences in the bacterial community could be found regarding the two reservoirs. *Bacteroidota* were significantly more abundant in Klingenberg, whereas *Planctomycetota* were significantly more abundant in Kleine Kinzig Reservoir (*p* < 0.001). This is consistent to results shown by Llirós et al. (2014) where the origin of the samples had the greatest effect on differences in the bacterial community composition, compared to season and water layer.

### 4.3. Proliferation as an autochthonous process in the reservoir

As fecal contamination of raw water is a major challenge for drinking water suppliers (Gunnarsdottir et al., 2020), an extremely important question for them as well as for health authorities is whether these observed high numbers of coliform bacteria are related to fecal contamination, which would possibly implicate health risks. To address this question, FIB *E. coli* and enterococci were monitored during the entire sampling period, as coliform bacteria do not permit correlation on fecal influence. Furthermore, the inflow into the reservoir was compared with the reservoir water. Additionally, the reservoir water was tested for fecal MST marker genes.

In Germany, drinking water reservoirs are surrounded by a drinking water protection zone in order to minimize hazards within the catchment area (see e.g. WHO, 2016). In addition, pre-reservoirs serve as a barrier, e.g. to reduce the direct introduction of storm water into the main reservoir, which can be detected by elevated numbers of microbial indicator parameters (Balzer et al., 2010; Kistemann et al., 2002). In contrast to coliform bacteria and HPC, the FIB did not reach high numbers during the time of mass proliferation, even more they were not detectable in the majority of the samples. Most of the enterococci were identified as *Enterococcus rotai*, a species known to occur mainly in invertebrates like mosquitoes (Hügler et al., 2014; Sedláček et al., 2013). In general, enterococci do not only occur in feces, but also in the environment (see Byappanahalli et al., 2012 and references therein). The concentration of *E. coli* – the bacterium proposed to be the best indicator for fecal contamination from human as well as livestock or wildlife sources (Edberg et al., 2000; Farnleitner et al., 2010) – was even lower than enterococci. RDA revealed no correlation between the proliferating coliform species and the measurement of FIB (Fig 6). Thus, the high numbers of coliform bacteria in the reservoirs were not accompanied with increased numbers of FIB. Therefore, it can be assumed that coliform growth is not due to fecal contamination.

The analysis of the Klingenberg inflow confirms this assumption. Klingenberg Reservoir (raw water and surface water) was compared with its main inflow (Wilde Weißeritz) (Fig. 8, Table S1). In 2018, the year with the mass proliferation of coliform bacteria, it could be observed that numbers of all three indicators were usually at least 10 times higher in the inflow than in the reservoir. Only during the summer months, i.e. during the mass proliferation, the numbers of coliform bacteria reached higher maxima in the reservoir than in the inflow. However, the fecal indicators *E. coli* and enterococci were significantly higher in the inflow (*p* ≤ 0.05). Furthermore, FIB were below detection limit (1 per 100 ml) in many samples in the reservoir, whereas in the inflow these bacteria were detectable in all samples. This shows that the reservoir and its inflow were significantly different. This and the fact that *E. asburiae* was not detected in the main inflow during the sampling supports the conclusion that mass proliferation of coliform bacteria is an autochthonous process within the reservoir.

**Fig. 8:**
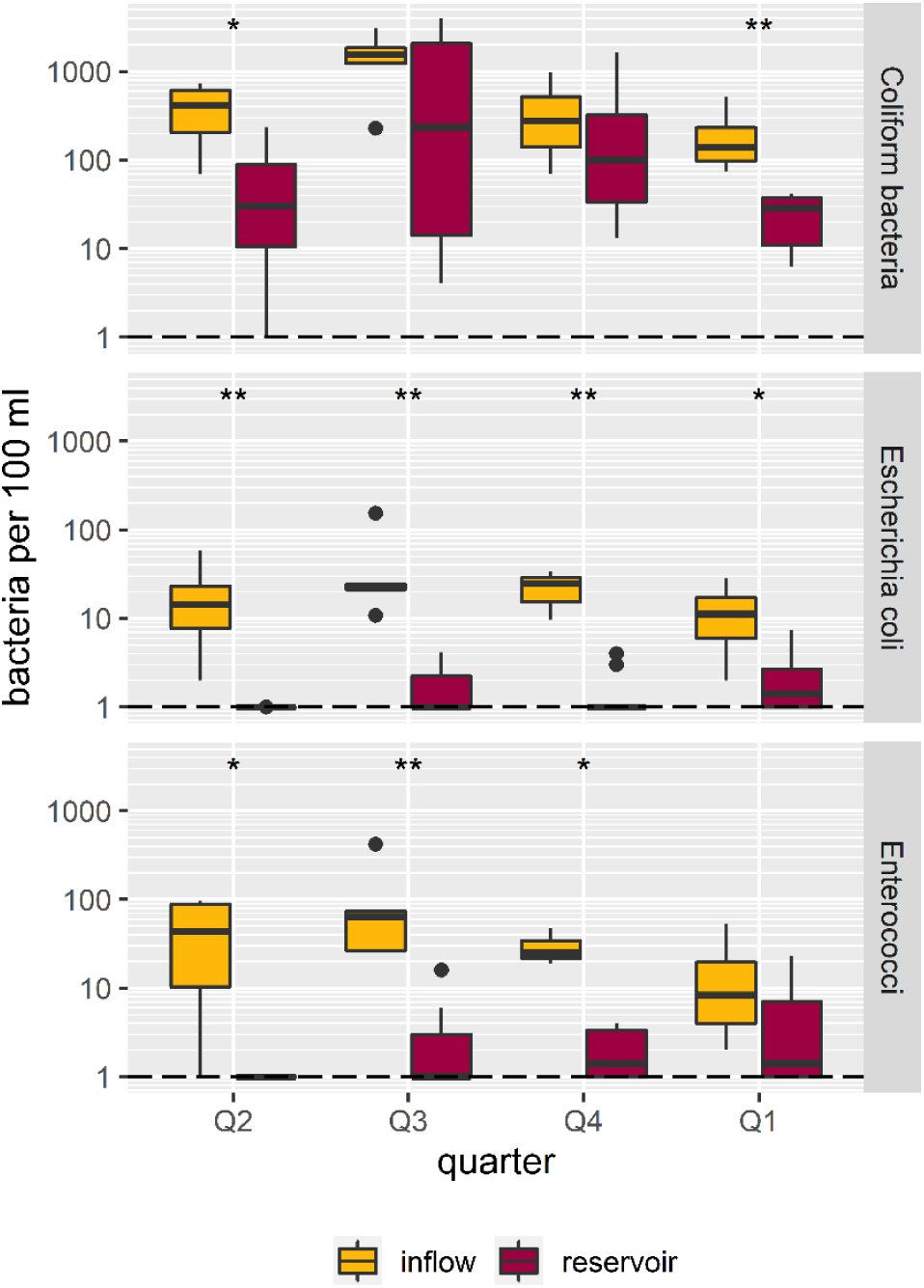
Differences in coliform bacteria, Escherichia coli and enterococci in the Klingenberg Reservoir compared to its inflow. Q2: Apr – June 2018, Q3: July – Sep 2018, Q4: Oct – Dec 2019, Q1: Jan – Mar 2019. Black line represents the detection limit of 1. For graphical illustration, values below detection limit (<1) were set to 0.99. For testing significant differences between the mean of inflow and reservoir samples the Mann-Whitney-U-test was performed (*: p ≤ 0.05, **: p ≤ 0.01).

For MST, specific *Bacteroidales* genes were tested during this study. These bacteria are one of the most abundant bacterial groups in the intestine of warm-blooded animals (Hagedorn et al., 2011). Therefore, quantification of *Bacteroides* 16S rRNA is a reliable method to estimate fecal contamination within water samples as well as the fecal origin, due to their co-evolution with specific hosts (Ahmed et al., 2016). During this study, unspecific *Bacteroides* 16S rRNA made around 0.2% of all 16S rRNA (2 x 10^3^ gene copies/ml). These numbers are relative low, as 1 g of feces in 1 L of water contains approximately 10^10^ gene copies of unspecific *Bacteroides* markers, which normally makes about one third of the fecal bacteria (Layton et al., 2006). Human- and ruminant-specific markers could not be detected. Furthermore, only one sample, namely the main inflow into the Klingenberg Reservoir, had a low fecal burden of pigs (4 x 10² gene copies/ml) in September. As this sampling spot is surrounded by forest, it is possible that the influence is caused by wild pigs, as they are also detected with this primer set (Lamendella et al., 2013). As this was the only sample with an influence from pig feces, it can be concluded that fecal contaminations in the inflow did not reach the reservoir. Fecal contamination results in an increase of *Bacteroides* markers, together with FIB (Stange and Tiehm, 2020). Furthermore, *Bacteroidales* are usually two to three orders of magnitude more abundant in human and animal intestine than coliform bacteria (Hagedorn et al., 2011). This was not the case in the reservoir waters. Thus, quantification of MST marker genes further suggested that the increase of coliform bacteria in the reservoirs did not correspond with fecal contaminations.

Together with the fact that the proliferating coliform species are rarely found in clinical material (Sanders and Sanders, 1997), and no *Firmicutes* could be detected in the 16S rRNA amplicon analysis of the reservoir waters, the aforementioned results indicate, that the growth of coliform bacteria within the reservoir is an autochthonous effect and is not due to fecal contamination.

### 4.4. Parameters affecting the proliferation of coliform bacteria

As fecal contamination is not the reason for the proliferation of coliform bacteria in drinking water reservoirs, the question arises what other factors possibly influence this process? Further questions are: Where, i.e. in which water depth or layer does the proliferation of coliform bacteria start? And how can these bacteria spread throughout the entire water body of the reservoirs? Therefore, RDA was conducted, to define parameters affecting the proliferation of coliform bacteria (Fig. 6).

For both proliferating species, *Enterobacter* spp. as well as *Lelliottia* spp., temperature was one of the mayor factors correlating with their occurrence, with *Lelliottia* proliferating earlier in summer compared to *Enterobacter*. One reason for that could be a lower growth temperature, since *Lelliottia* spp. have been shown to propagate even at lower temperatures than *Enterobacter* spp. (Izard et al., 1981; Leclerc et al., 2001). The connection of seasonal variability of coliform bacteria and temperature in surface waters, river and soil has been reported in several cases before (Cho et al., 2016; Hong et al., 2010; Jeon et al., 2019; LeChevallier et al., 1996). Higher water temperatures as well as water scarcity are predicted to increase in many regions of the world due to climate change (Bates et al., 2008; Dokulil et al., 2006; Mosley, 2015). In the last 40 years, water temperature in lakes has increased around 0.1 to 1.5 °C (Bates et al., 2008). Furthermore, stratification period has lengthened (Bates et al., 2008). This affects many physicochemical as well as biological parameters (Delpla et al., 2009), and we anticipate that the mass proliferation of coliform bacteria occurs even more frequently in future.

But not only water temperature, also oxygen content as well as nutrients and metals are important for bacterial growth. RDA also revealed oxygen and manganese as factors explaining the variance of coliform species in the reservoirs. As a consequence of stratification in reservoirs, oxygen consumption in the hypolimnion can lead to the solution of metals and nutrients from the sediment. Highest release of manganese from sediments into the water occurs in summer, when reservoirs are stratified (Munger et al., 2019). As facultative anaerobic bacteria coliform bacteria may have an advantage compared to other bacteria in this environment. Furthermore, in Klingenberg Reservoir correlation with total phosphate was found. *Enterobacter* spp. are known as plant growth promoting bacteria (Andrés-Barrao et al., 2017; Oh et al., 2018; Taghavi et al., 2010) capable of phosphate mobilization (Gyaneshwar et al., 1999; Mezzatesta et al., 2012; Vazquez et al., 2000). This could explain the correlation of coliform bacteria and total phosphate.

Where does the mass proliferation begin? It is conceivable that mass proliferation could start near the thermocline, as inactivation due to UV-light is lower in this region (Davies-Colley et al., 1994). Additionally, particles together with associated bacteria settle down, as well as nutrients form an upward flux (Davis et al., 2005). It is therefore possible that this layer is enriched with particles of dead algae, which can serve as nutrients for heterotrophic bacteria like the coliform bacteria (Cole, 1982; Kouzuma and Watanabe, 2015; McFeters et al., 1978), thus leading to increased concentrations of bacteria near the thermocline (Davis et al., 2005; McDonough et al., 1986).

Another open question is how the coliform bacteria manage to colonize the entire reservoir in all depth layers, even when the reservoir is stratified. Due to drinking water production, raw water is extracted from the hypolimnion of the reservoirs. This withdrawal influences both volume as well as temperature of the hypolimnion (Çalışkan and Elçi, 2009; Casamitjana et al., 2003; Mi et al., 2019; Weber et al., 2017). As the hypolimnion warms up, thermal stability decreases. Additionally, withdrawal can produce buoyancy forces and the withdrawal layer may even intersect the thermocline (Çalışkan and Elçi, 2009; Fischer et al., 1979). This could explain how the bacteria may overcome the thermocline and occur in all depth layers at least in smaller reservoirs. Furthermore, *E. asburiae* is reported as a quorum sensing bacterium, capable to communicate with surrounding bacteria (Lau et al., 2013; Lau et al., 2020). However, the adaption of these bacteria to their productivity-poor environment is not yet solved. These habitats pose a challenge for bacteria, especially when it comes not only to survive but also to proliferation (Kundu et al., 2020). Therefore, further research is needed, to understand the exact place of the origin of proliferation, the ways the bacteria distribute in the water column and the adaptions of these bacteria to their environment.

The mass proliferation in the two reservoirs seems to be caused by different factors, as not only RDA, but also the time of mass proliferation determines. In Klingenberg Reservoir it is starting much later, in the end of summer, compared to Kleine Kinzig Reservoir in early summer season. Furthermore, *Lelliottia* spp. is detectable in both reservoirs during the whole year, with low numbers between January and May. *Enterobacter*, however, was not present in the reservoir, only when it comes to mass proliferation. Furthermore, highest numbers in Klingenberg Reservoir have been observed in raw water depth, in Kleine Kinzig Reservoir, however, the opposite was the case and highest numbers were measured in the surface layer. Differences between the two reservoirs were observed regarding temperature, as well as nutrient content. Especially in Klingenberg, nitrate, silicate and TOC are higher. Taken together, this leads to the conclusion that both effects are influenced by different factors. Water temperature might be one of the leading causes of mass proliferation of coliform bacteria. Yet, additional factors seem to be important (e.g. manganese, oxygen, phosphate) that have to be investigated in further research.

### 4.5. Implications for drinking water treatment

Climate change leads to enhanced water temperatures and extreme water events. So droughts, heavy rain events and flooding are expected to happen more frequently. Thus, it is anticipated, that mass proliferations of coliform bacteria in drinking water reservoirs also occur more often in the near future. At some reservoirs, these events are observed almost on a yearly basis. This challenges drinking water treatment. Currently, water treatment plants treating reservoir water mainly use conventional treatment techniques like flocculation and filtration, followed by disinfection. Even with a fully functional treatment technology, it is conceivable that coliform bacteria might overcome drinking water treatment in times with high densities of more than 10^4^ bacteria per 100 ml in raw water. If coliform bacteria reach drinking water distribution systems, they might proliferate in biofilm or in sediments within the system (Camper, 1993; Edberg et al., 1994; LeChevallier et al., 1996). This could lead to the exceeding of limit values (e.g. according to EU drinking water directive, coliform bacteria should not be detectable in 100 ml), often followed by countermeasures like additional disinfection measures or even boiling advices.

Thus, it is essential to increase our knowledge about coliform bacteria, especially with regard to strains capable to propagate in drinking water reservoirs and potentially also in drinking water systems under certain conditions. Genome analyses in combination with specific growth experiments are conceivable to answer open questions with regard to the hygienic relevance of these bacteria and their adaptations to oligotrophic environments. Furthermore, monitoring of proliferation events on a high spatial and temporal resolution could give further insights and answer remaining questions.

## 5. Conclusion

- In our monitoring of drinking water reservoirs, we detected seasonal differences in numbers of coliform bacteria.
- During summer months, the maximum density of coliform bacteria reached values of more than 10^4^ bacteria per 100 ml in these oligotrophic environments, not only in the surface but also in raw water (used for drinking water production). This represents an increase of 4 orders of magnitude compared to the winter season.
- At this time, only one or two strains of coliform bacteria dominated the entire water body of the reservoir, belonging to the genera *Lelliottia* or *Enterobacter*. Closely related strains were found in different reservoirs all over Germany.
- Our study demonstrates that proliferation was not due to fecal contamination but is an autochthonic process within the water column of the reservoirs.
- We conclude that this sudden increase of single species can be considered as a mass proliferation of coliform bacteria or “coliform bloom” within the reservoir.
- Microbial community was dominated by *Actinobacteriota*, *Bacteroidota* and *Proteobacteria* and did not change significantly during periods of mass proliferation. *Enterobacterales* only make around 0.04% of the community.
- RDA revealed correlation of proliferating coliform species with increased water temperatures, lower oxygen content as well as nutrients and metals (phosphate, manganese).

## Acknowledgment

This research was funded by the German Federal Ministry of Education and Research (BMBF grant no. 02WGR1426A-G, project TRUST) and the German Technical and Scientific Association for Gas and Water (DVGW grants no. W 201720 and W 201823). We thank Margret Sommer and Gerhard Biwer from Kleine Kinzig Reservoir, as well as Thilo Hegewald from Klingenberg Reservoir for providing samples and access to the reservoirs. We further thank the water suppliers Fernwasserversorgung Elbaue-Ostharz, Harzwasserwerke, Stadtwerk am See and Wasserverband Siegen-Wittgenstein for providing water samples. Furthermore, we thank Monika Bösl, Lydia Grabner, Luisa Telpl and Carolin Schweikart (TZW Karlsruhe), as well as Jana Angermann, Annett Schröter, Claudia Kupfer and Susann Bromberger (TZW Dresden) for laboratory support at TZW.

